# Temporal variability in host availability alters the outcome of competition between two parasitoid species

**DOI:** 10.1101/2024.01.30.577865

**Authors:** Hua Wang, Tiantian Liu, Shucun Sun, Owen T. Lewis, Xinqiang Xi

## Abstract

1. Variability in the availability of resources through time is a common attribute in trophic interactions, but its effects on the fitness of different consumer species and on interspecific competition between them are not clearly understood.
2. To investigate this, we allowed two parasitoid species, *Trichopria drosophilae* and *Pachycrepoideus vindemiae*, to exploit *Drosophila* host pupae under three temporal variability treatments, either on their own or simultaneously.
3. When tested individually (in the absence of interspecific competition), both parasitoid species had lower fitness when hosts were exposed for a short duration at high density than when exposed for a long duration at low density. When both parasitoid species exploited hosts simultaneously, interspecific competition significantly decreased the number of offspring for both parasitoid species. The outcome of this interspecific competition depended on host temporal variability, with *T. drosophilae* or *P. vindemiae* dominating in short and long host exposure treatments, respectively.
4. These results can be explained by the combination of host availability and egg load of female adult parasitoids. When abundant hosts are provided for a short period, the ample mature eggs of the proovigenic *T. drosophilae* enable them to exploit hosts more efficiently than *P. vindemiae*, which is synovigenic. However, *P. vindemiae* is an intrinsically superior competitor and dominates when multiparasitism occurs. Multiparasitism is more frequent when hosts are at low levels relative to the egg load of the parasitoids.
5. Our results clearly demonstrate that resource temporal availability can alter the outcome of competition between consumers with different reproductive traits.

## Introduction

Variability through time in the availability of resources is a common attribute in many ecosystems (Ostfeld & Keesing 2000). Abrupt, ephemeral changes in resource availability, such as periodical cicada outbreaks in north America, play a significant role in regulating population and community dynamics of consumers (Yang et al., 2008). Less extreme temporal variations in most abiotic and biotic resources are widespread (Mirth, Saunders & Amourda 2021), and are expected to have marked effects on consumer species. Generalist consumers, especially those that have long lifespans, can maintain whole-life fitness in the face of resource temporal variation through behavioral adaptations such as prey switching (Setälä et al., 2022). However, the performance of specialist consumers with low mobility and short lifespans may suffer considerably during short periods of low resources, even if longer-term total resource availability is unchanged (Yang et al., 2010). Previous studies demonstrate that phenological match between consumer and resources determines resource use efficiency, but it is less clear how temporal variation in resources changes the dynamics of competing consumers sharing the same resources (Lopez et al. 2017; Ceron et al. 2022). Increasing our understanding of consumer-resource temporal dynamics is particularly important since ongoing environmental change is not only altering the abundances of resource and consumer species, but also changing their temporal variability (Bale et al. 2002).

Parasitoids are diverse, ubiquitous and important enemies of insects and other arthropods (Godfray 1994). Most parasitoids have short lifespans as adults and are trophically specialized, with few (often closely related) host species which can often be exploited only at specific developmental stages. These characteristics make parasitoids especially susceptible to the spatial and temporal dynamics of their hosts (Hassell 2000). A variety of mechanisms can lead to considerable variation in host availability through time, including variation in the quantity and quality of host plant resources in the case of herbivorous insects (Hicks et al., 2007; Frank 2021) and the impact of abiotic factors that can lead to synchronized population peaks and discrete generations for multivoltine hosts (Lima, Stenseth & Jaksic 2002; Greiser et al., 2022). For example, after extremely cold winters individuals of the multivoltine moth *Adoxophyes honmai* end dormancy within a narrow window, so that their offspring occur as discrete generations with intervals of ∼30 days. In this case, resources for parasitoids have low temporal homogeneity but are at high density when available (Bjørnstad, Nelson & Tobin 2016). In contrast, after warm winters, dormancy tends to end asynchronously so that offspring of different generations overlap to a high degree (Bjørnstad, Nelson & Tobin 2016), resulting in temporally-homogeneous but low-density resources for their parasitoids.

Temporal heterogeneity in host availability will alter the host use efficiency of parasitoids (Cusumano et al., 2022), and is expected to have differing effects depending on parasitoid life history strategy. Proovigenic parasitoids have eggs that are mature at or shortly after adult emergence and lay their eggs over a relatively short period, while synovigenic parasitoids have eggs that mature gradually during their longer lifespan (Jervis et al., 2001; Segoli & Rosenheim 2013a). Reproductive success of parasitoids may be limited by the availability of mature eggs for synovigenic parasitoids, and by the availability of suitable hosts for proovigenic parasitoids (Rosenheim 1996; Segoli et al., 2018). We predict that when small numbers of hosts are continuously exposed to parasitoids over an extended period, long-lived synovigenic parasitoids will be favoured, but when hosts occur at high density for short periods, proovigenic parasitoids with plentiful mature eggs will be better able to exploit host resources. Furthermore, high intraspecific competition associated with superparasitism is expected when hosts are few relative to parasitoid egg loads, decreasing the fitness of parasitoids, especially solitary proovigenic parasitoids (Santolamazza-Carbone & Rivera 2003; Duvall et al., 2018).

When the same host species is shared by more than one parasitoid species, these parasitoids can compete both extrinsically and intrinsically (i.e., as larvae within individual hosts, Harvey, Poelman & Tanaka 2013; Ode, Vyas & Harvey 2022). If hosts are available to parasitoids at low density over a long duration, higher rates of multiparasitism will occur, increasing the strength of intrinsic interspecific competition. In contrast, when host availability is more heterogeneous through time, extrinsic competition driven by host use efficiency will dominate (Hassell, 2000). Differences in the life-history characteristics of parasitoids will therefore influence the circumstances when they can achieve competitive superiority. For example, synovigenic parasitoids have few eggs but invest more in each, with the ovipositing wasps injecting venom to kill competing larvae in multiparasitized hosts (Wang et al., 2016). Therefore, host temporal availability will indirectly determine the fitness and ultimately the abundance of competing parasitoid species by changing the relative importance of extrinsic and intrinsic processes in interspecific competition (Segoli & Rosenheim 2013b).

Here, we investigate experimentally the effect of host temporal variability on parasitoid fitness and competitive outcomes in a model host-parasitoid system comprising *Drosophila melanogaster* (Diptera: Drosophilidae) and two of its pupal parasitoids, *Trichopria drosophilae* and *Pachycrepoideus vindemiae*. We allow the parasitoids to attack the shared host species, either alone or in competition, and under one of three temporal variability treatments. Our results suggest an important role for temporal resource availability in modulating fitness and the outcome of interspecific competition in host-parasitoid systems.

## 2 Materials and methods

### 2.1 Study system

*Drosophila* flies are distributed widely across the world (Markow 2015). These insects are a biologically-relevant model for understanding the consequences of fluctuations in resource availability for host-parasitoid dynamics because their larval resources, such as rotten fruits or fungi, are typically ephemeral and show pronounced fluctuations in availability; for example, fungal fruiting bodies often occur as short-lived pulses after heavy rain (Dulay et al., 2020). *Drosophila* fly populations track these changes, increasing rapidly when resources become available and decreasing abruptly once they are depleted (Breitmeyer & Markow, 1998). *Drosophila* species are hosts for more than 100 parasitoid species (Lue et al., 2021) worldwide. *Trichopria drosophilae* and *Pachycrepoideus vindemiae* are cosmopolitan pupal parasitoids of Drosophilidae with wide host ranges. They are easily maintained in captive culture and have been widely studied in investigations of host-parasitoid dynamics and biocontrol (e.g., Wang et al., 2021).

### 2.2 Insect rearing

Laboratory colonies of *D. melanogaster*, *T. drosophilae* and *P. vindemiae* were established using insects collected from Zhonggong Grape Experimental Station, Shandong Province, China in 2019. They were cultured under laboratory conditions (25 ± 1L, L:D = 16:8, RH: 60∼70%) using a cornmeal-based artificial diet composed of yeast, agar, and sucrose. In preparation for the experiments we placed six standard 100 ml plastic bottles with 20ml of artificial diet into a plastic box (length × height × width = 40 × 20 × 20 cm) with the opening covered in gauze, and then released ∼40 pairs (♀:♂=1:1) of adult flies into the box to lay eggs in the bottles for 24 hours. To obtain an abundance of fresh pupae, we replaced the old medium in the box with two bottles of fresh medium every 7 days. After fly larvae pupated, 40 1 to 2 day old fresh pupae were transferred into vials (diameter = 2.5 cm, height = 9.5 cm), and four pairs (♀:♂ = 1:1) of adult parasitoids were introduced and allowed to lay eggs for 24 hours. Parasitoids were used on the day of emergence in the subsequent experiments, and adult wasps were fed 10% sucrose water solution. To continuously obtain experimental parasitoids, a batch of parasitoids was expanded every 7 days.

### 2.3 Effect of the host availability on parasitoid fitness

To test the effects of temporal variability in host resource availability on parasitoid performance, we released a pair (♀:♂ = 1:1) of *T*. *drosophilae* or *P. vindemiae* into a vial containing 8, 20 or 40 fresh *D. melanogaster* pupae and allowed parasitoids to lay eggs for 24 hours. This procedure was repeated for each pair of parasitoids transferred into a vial with the appropriate number of fresh host pupae every 24 hours, for a duration of 10, 4 and 2 days, respectively. Therefore, total resource availability was held constant across the experiment (80 pupa-days) but with three temporal variability treatments: 8 pupae/day for 10 days, 20 pupae/day for 4 days, or 40 pupae/day for 2 days. Female parasitoids were allowed to feed *ad libidum* on 10% sucrose water solution. Parasitoids that did not lay eggs within 10 minutes of being introduced to the vials were discarded and the corresponding vials were not included in the analyses. Twenty replicates were set up for each parasitoid species for each of the three host temporal variability treatments.

To assess the host-use efficiency of parasitoids, 24 hours after parasitoids were removed from the vials we dissected all fly pupae in ten replicates of each treatment under a stereomicroscope to search for parasitoid larvae. Superparasitism was identified when two or more parasitoid larvae were found in a single pupa. When *T*. *drosophilae* larvae reach the late first instar (approximately 72 hours after eggs are laid) and have a well-developed caudal appendage and strong mandibles, cannibalism can occur (Wang et al., 2016), leaving behind traces of cannibalized larvae. No cannibalism was found in our experiment. In the remaining 10 replicates of each treatment we recorded the number of offspring of female *T*. *drosophilae* or *P. vindemiae*.

Replicates where there were no parasitoid progeny as a result of females not laying eggs, or where only male progeny emerged as a result of females not having mated, were not included in analyses, because parasitoid oviposition behavior may change according to mating status. Consequently, the final number of replicates varied between 5 and 10 per treatment (see Results for detailed information).

### 2.4 Effect of the host availability on parasitoid inter-specific competition

To investigate whether temporal variation in host availability can shift the outcome of inter-specific competition between two parasitoid species, we released a mated female *T. drosophilae* and a mated female *P. vindemiae* simultaneously into vials with the temporal variability treatments described in section 2.2. Twenty replications were set up for each of the three host temporal variability treatments.

To quantify extrinsic competition between the two parasitoid species, we again dissected fly pupae in ten replicates of each treatment 48 hours after releasing the adult parasitoids. Superparasitism was identified when two or more larvae from the same parasitoid species were found in the same host pupa, and multiparasitism was identified when we found larvae of both parasitoid species together. Eggs and larvae of the two parasitoid species could be distinguished easily based on differences in morphology. The remaining vials were used to record the number of offspring of *T. drosophilae* or *P. vindemiae*. Vials where no parasitoids or only male parasitoids emerged were not included in further analyses. Final numbers of replicates of each treatment were 9∼10.

### 2.5 Statistical analyses

All analyses used the statistical software R version 4.2.1 (R Core Team 2022). We employed generalized linear models (GLMs) with Poisson error distribution to test if the number of emerged parasitoids, number of parasitized pupae and the occurrence of superparasitism or multiparasitism were significantly different among the three host temporal variability treatments. To assess the significance of individual factor levels within GLMs we employed Tukey post-hoc tests using the “*glht*” function in the *multcomp* package (Bretz, Hothorn & Westfall 2016).

## 3 Results

### 3.1 Effect of host temporal variability on parasitoid fitness

In the single-species experiments, host availability has a significant effect on the number of emerged offspring of both *T*. *drosophilae* (*z* = −2.10, *p* =0.035) and *P. vindemiae* (*z* = 13.88, *p* < 0.001). Specifically, the number of emerged *P. vindemiae* offspring was highest when 8 pupae/day were exposed for a total of 10 days (Figure 2b), while the number of *T*. *drosophilae* offspring was highest when 8 pupae/day were exposed for 10 days, or 20 pupae/day for 4 days (Figure 2a).

**Figure 1.**
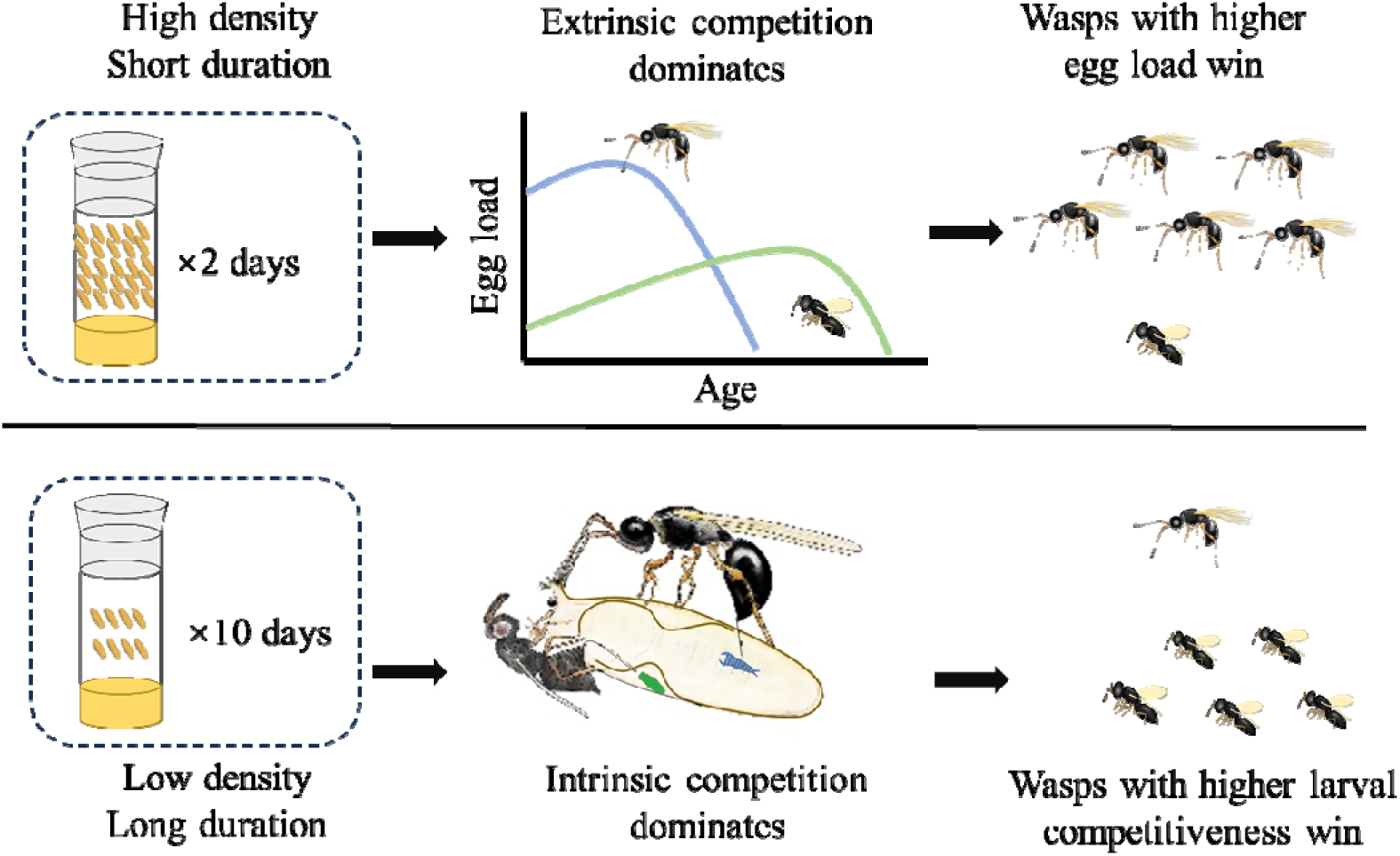
The impact of host temporal variability on interspecific competition between two parasitoid species.

**Figure 2.**
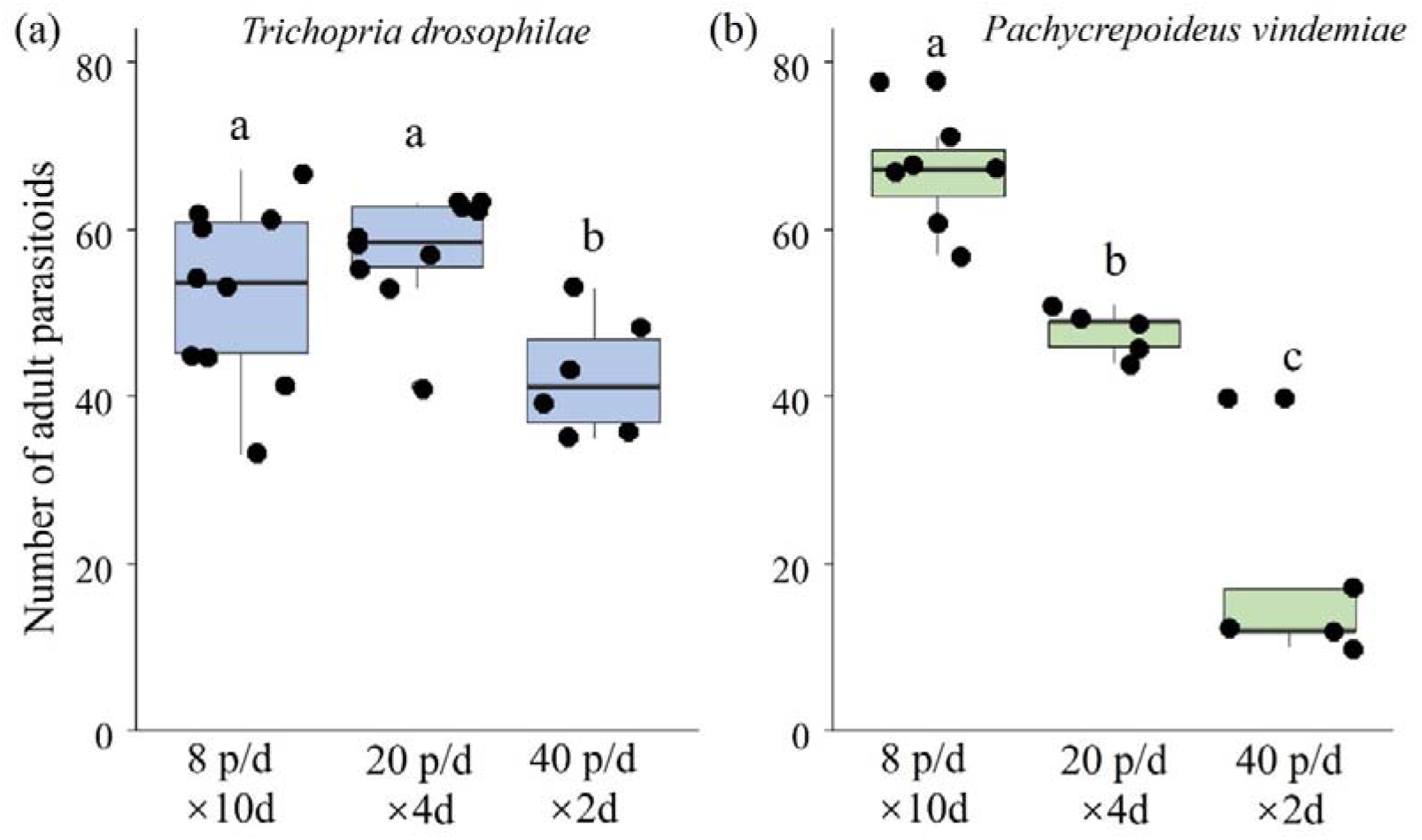
Numbers of adult *Trichopria drosophilae* or *Pachycrepoideus vindemiae* that emerged when host pupae were exposed to parasitoids separately. Data points are shown around the boxes. The solid circles represent the data points. Different letters above the boxes indicate significant differences among treatments (*p* < 0.05).

Dissecting *Drosophila* fly pupae exposed to *T. drosophilae* or *P. vindemiae* showed that the total number of pupae parasitized by *T. drosophilae* tended to differ among treatments (*z* = 1.89, *p* = 0.059, marginally non-significant), with more pupae parasitized by *T. drosophilae* when 8 pupae/day were available for 10 days, or when 20 pupae/day were available for 4 days. Additionally, incidence of superparasitism (host pupae with more than one parasitoid larva) was significantly higher in these two treatments than in the lowest temporal variability treatment (*z* = −8.76, *p* < 0.001, Figure 3a). In contrast, the number of pupae parasitized by *P. vindemiae* decreased as host temporal variability increased (*z* = −10.27, *p* < 0.001). Only one pupa with two larvae of *P. vindemiae* was recorded (Figure 3b).

**Figure 3.**
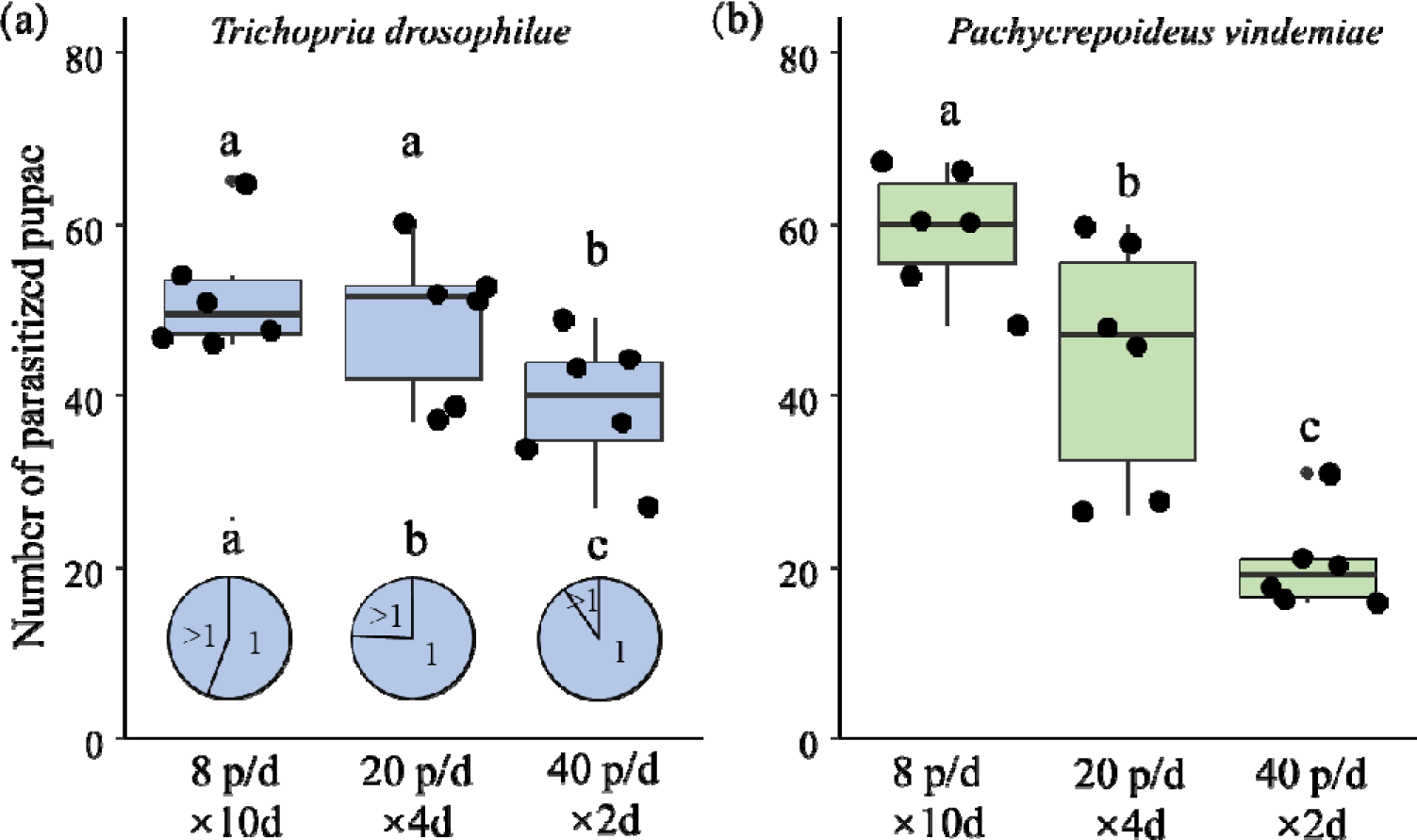
Numbers of pupae parasitized by *Trichopria drosophilae* or *Pachycrepoideus vindemiae* when host pupae were exposed to parasitoids separately. Data points are shown around the boxes. The pie chart marked with “1” and “>1” indicates the proportion of parasitized pupae with one or more than one parasitoid larva, respectively. Different letters above the pie charts and boxes indicate significant differences among treatments (*p* < 0.05).

### 3.2 Effects of host temporal variability on parasitoid competition

When hosts were exposed to the two parasitoid species simultaneously, host temporal variability changed the outcome of interspecific competition (*z* = 4.87, *p* < 0.001). Specifically, significantly more *P. vindemiae* than *T. drosophilae* were found when they were provided with 8 pupae/day for 10 days (*z* = 9.776, *p* < 0.001, Figure 4) but significantly more *T. drosophilae* than *P. vindemiae* were found when 40 pupae/day were available for 2 days (*z* = 3.222, *p* = 0.001). Numbers of *P. vindemiae* and *T. drosophilae* did not differ significantly when 20 pupae/day were available for 4 days (*z* = −0.864, *p* = 0.388).

**Figure 4.**
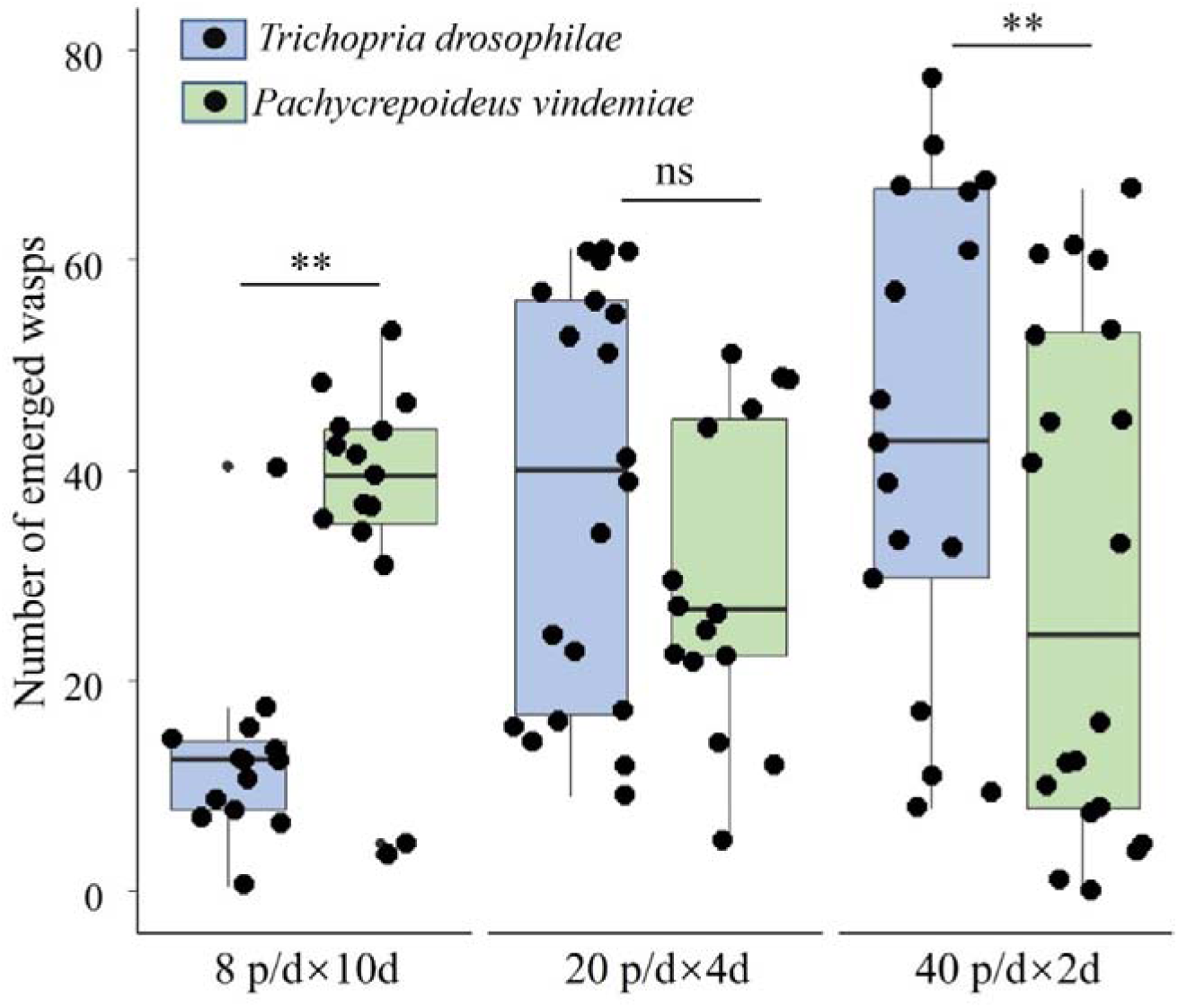
Numbers of adult *Trichopria drosophilae* and *Pachycrepoideus vindemiae* emerged when host pupae were simultaneously provided. Boxplots represent the median, the quartiles, and the maximum and minimum values, which generated from data including the extreme outliers. Data points are shown around the boxes. “**” and “ns” above boxes indicate significant (*p* < 0.05) and non-significant difference between the two parasitoids, respectively.

Dissection revealed that when hosts were exposed to the two parasitoid species simultaneously, the number of pupae parasitized by *T. drosophilae* increased significantly as host variability increase (*z* = 4.458, *p* < 0.001, Figure 5), but the number of pupae parasitized by *P. vindemiae* decreased significantly with host variability (*z* = −6.253, *p* < 0.001). In addition, the number of pupae parasitized by both *T. drosophilae* and *P. vindemiae* decreased significantly as host variability increased (*z* = −8.576, *p* < 0.001). The number of pupae with more than one *T. drosophilae* or *P. vindemiae* larva decreased with increasing host variability (*T*. *drosophilae*: *z* = −3.172, *p* = 0.002, *P. vindemiae*: *z* = −3.974, *p* < 0.001).

**Figure 5.**
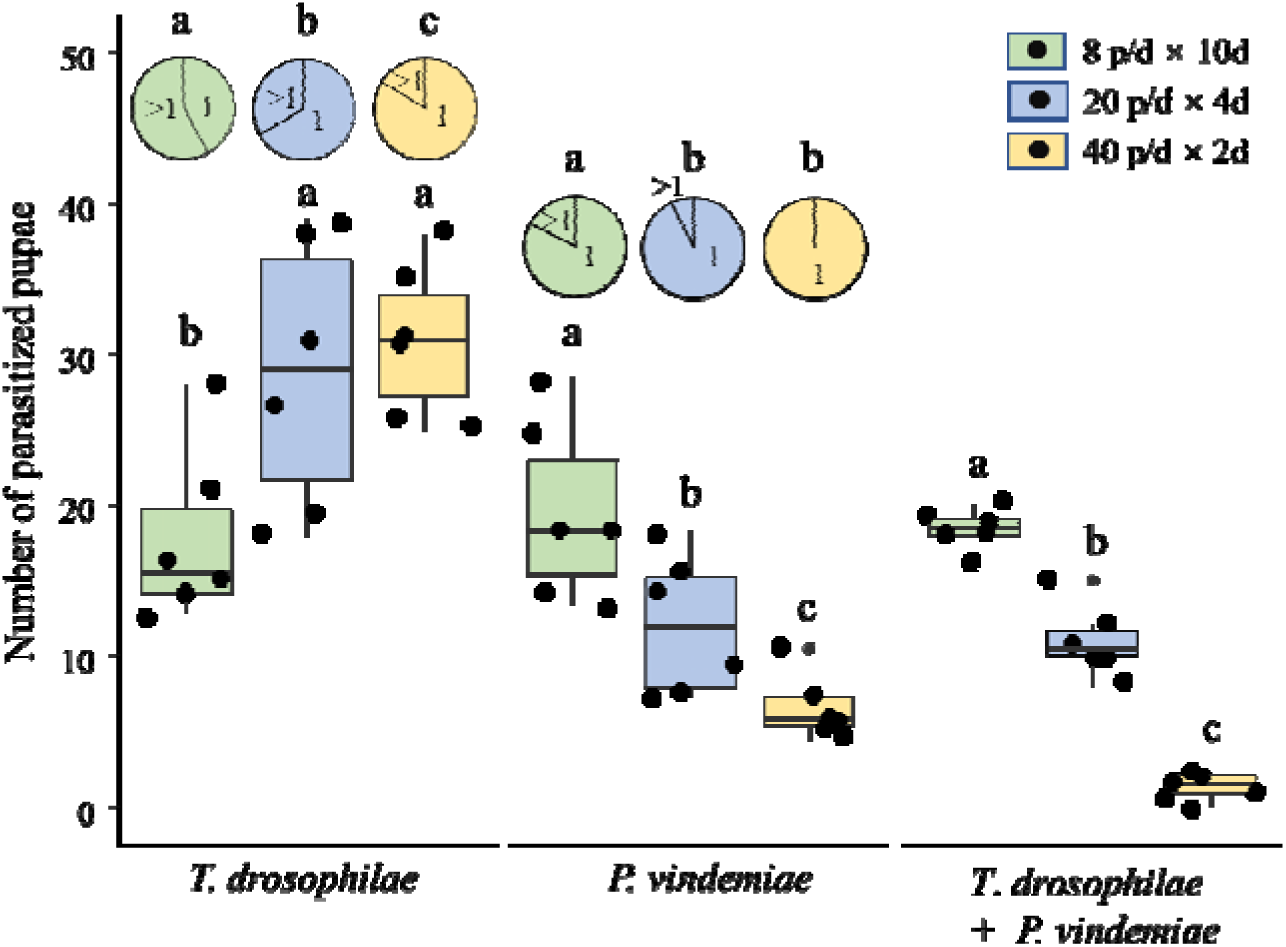
Numbers of pupae parasitized by *Trichopria drosophilae* and *Pachycrepoideus vindemiae* when host pupae were exposed simultaneously. Boxplots represent the median, the quartiles, and the maximum and minimum values, which were generated from data including the extreme outliers. Data points are shown around the boxes. The pie chart marked with “1” and “>1” indicates the proportion of parasitized pupae with one or more than one parasitoid larva, respectively. Different letters above the pie charts and boxes indicate significant differences among treatments (*p* < 0.05).

## Discussion

Several previous studies have demonstrated that ephemeral resource pulses have pronounced effects on consumer fitness and ecosystem function (Holt 2008; Ostfeld & Keesing 2000; Yang et al., 2008; Yang et al., 2010). In this study we found that temporal variability in host resource availability has similarly marked effects on parasitoid fitness and interspecific competition. Compared to free-living predators, the fitness of parasitoids is more closely tied to host resource variability, since it determines not only the strength of competition among female parasitoids for hosts, but also the intensity of intrinsic competition for parasitoid larvae within hosts. By adding experimental evidence involving short-lived consumers, our study broadens knowledge of how temporal resource variability mediates consumer performance.

Temporal host availability affects the fitness of *T. drosophilae* and *P. vindemiae* differently. A principal explanation for this is contrasting patterns of egg maturation in the two parasitoid species. Newly-emerged *T. drosophilae* have approximately 50 oocytes, almost six times as many as *P. vindemiae* (Table S2). Furthermore, 38% of eggs in *T. drosophilae* are already mature when females hatch from their pupae, while oocytes mature slowly with age in adult female *P. vindemiae* (Figure S1). As a result, female *T*. *drosophilae* laid more eggs and had more offspring than *P. vindemia*e when 40 pupae/day were supplied for 2 days. Therefore, parasitoid fitness is dependent, to some extent, on the temporal aggregation of hosts.

A further reason for the different effects of host temporal availability on fitness of the two species is that *T*. *drosophilae* is poorer at distinguishing unparasitized pupae than *P. vindemiae* (Wang et al., 2016). A higher frequency of superparasitism was found for *T*. *drosophilae* when 8 host pupae/day were provided for 10 days, leading to wasted eggs and higher larval mortality, since each *Drosophila* pupa can support the development of only one parasitoid. Reproduction of *P. vindemiae* is limited by the length of the egg-laying period, and they had higher host use efficiency and laid more eggs when hosts were provided over an extended period.

Proovigenic and synovigenic have been assumed to represent a fundamental dichotomy in parasitoid egg maturation strategies, but Ellers & Jervis (2004) found that only 1.8% of surveyed species were strict proovigenic. Likewise, proovigenic parasitoids constitute a much smaller proportion of parasitoids of *Drosophila* flies in natural ecosystems than synovigenic parasitoids (Fleury et al. 2009). This could be because the high energy costs of maintaining mature eggs mean that proovigenic life histories come at the expense of shortened lifespans and reduced host efficiency when the temporal availability of host resources is highly unpredictable (Segoli et al. 2018; Rosenheim et al. 2008). The abundance of *Drosophila* flies in natural ecosystems is closely dependent on resources such as rotten fruits or mushroom that themselves have high temporal variability, further influencing the temporal availability of hosts for parasitoids. When *Drosophila* flies are present at low density over a long period, or phenological mistmatch occurs between flies and wasps, the fitness of proovigenic parasitoids will be greatly reduced.

In addition to its differential effects on parasitoid fitness, we found that host temporal variability also shifted the outcome of inter-specific competition between parasitoids, and their resulting population dynamics. Inter-specific competition between parasitoids comprises two closely linked processes, extrinsic and intrinsic competition (Harvey, Poelman & Tanaka 2013; Ode, Vyas & Harvey 2022). In fact, *T. drosophilae* and *P. vindemiae* spend similar time in handling the host pupae (Table S1), but the higher egg load of *T. drosophilae* enables it to attack more host pupae successfully than *P. vindemiae* when abundant pupae are available to both species simultaneously over a short time period. Furthermore, both parasitoid species tend to avoid ovipositing in pupae that have been parasitized by competitors, as indicated by the much lower frequency of hosts containing eggs of both parasitoid species when temporal variability in host availability was high. Therefore, ephemeral bursts of host availability will tend to increase the superiority of adult *T. drosophilae* in extrinsic competition.

The outcome of interspecific competition was reversed when hosts were exposed at low density over a long period. Under these conditions, more adult *P. vindemiae* than *T. drosophilae* emerged from hosts, even though both parasitoid species parasitized similar numbers of pupae (indicating similar extrinsic competitive ability). This can be explained largely by the inferiority in inter-specific competition and higher frequency of intra-specific intrinsic competition of *T. drosophilae*. A higher frequency of multiparasitism was found in the low-host variability treatment which tend to increase the competitive advantage of *P. vindemiae*. Several physiological and behavioural characteristics may contribute to the competitive superiority of larval *P. vindemiae*. First, larvae of *P. vindemiae* have lower nutrient requirements than those of *T. drosophilae*, with higher survival when competition reduces nutrient availability in host pupae (Wang et al., 2021). Second, the competitive ability of *P. vindemiae* is likely to be enhanced by the ability of adult females to inject venom into the host at the time of oviposition, which can kill larval competitors (Wang et al., 2016). Finally, larval *P. vindemiae* live as ectoparasitoids outside the host pupa and can even take arval *T. drosophilae* that live within the pupae as host, since *P. vindemiae* is facultative hyperparasitoid (Van Alphen & Thunnissen, 1982; Wang & Messing, 2004).

Parasitoids are among the most diverse multicellular taxa in the world (Godfray 1994; Forbes et al., 2018), and their hosts are typically scarce and unevenly distributed. Thus, partitioning host resources in time or space is thought to be necessary to inhibit competitive exclusion between parasitoids sharing the same hosts (Murdoch, Briggs & Nisbet 1996; Bonsall, Hassell & Asefa 2002). Trade-offs between dispersal and competitive ability may also help competing parasitoids coexist on discrete resource patches (Amarasekare 2000; Hackett-Jones, Cobbold & White 2009). In this study, we showed that *P. vindemiae* and *T. drosophilae* win interspecific competition at low and high host variability, respectively. Considering the fluctuating dynamics of host populations in natural ecosystems, temporal shifts in competitive outcomes may, to some extent, inhibit competitive exclusion and promote coexistence of parasitoids, even without significant niche partitioning.

## Acknowledgment

We thank Shandong Academy of Agricultural Science for providing studied flies and wasps. Zhonghui Qiu helped dissect *Drosophila* pupae. This study was financially supported by the National Science Foundation of China (32022409).

