## Supplemental for "Temporal variability in host availability alters the outcome of competition between two parasitoid species"

1 **Materials and methods**

**1.1 Oviposition dynamic of two parasitoids**

To observe the daily egg maturation and lifetime fecundity, and calculate the ovigeny index of *T. drosophilae* and *P. vindemiae*, we introduced a newly emerged mated female of each species into a vial with 50 fresh host pupae, which is more than maximum mature eggs for the two studied wasp species. Females that did not lay eggs within 10 minutes after being introduced into vials were discarded. We removed adult parasitoids from vials after 8h and transferred them into another vial with 10% sucrose water solution for 24h to allow eggs to mature. The number of mature eggs each day for each parasitoid species was estimated from the number of emerged parasitoids. We dissected the parasitoids 15 days later to count the number of mature eggs left in their ovaries. The ovigeny index is calculated by dividing the number of mature eggs present at emergence by the lifetime production of eggs of a female parasitoid (Jervis et al., 2001)

1.2 **Reproductive traits of two parasitoids**

**Host handling time**

To estimate the host use efficiency of *T. drosophilae* and *P. vindemiae*, we recorded drumming, drilling and oviposition time (time needed for a wasp to complete an oviposition bout) of eight and nine newly mated females of each species individually. The total host handling time was estimated as the sum of the drumming, drilling and oviposition steps.

**Oocyte number and size**

Newly emerged parasitoid females were randomly selected to investigate the traits of eggs. Female parasitoids were first killed at -20 °C and then dissected in a petri dish with a drop of phosphate buffered saline (PBS). The length of the three to five largest oocytes and the total number of mature oocytes per individual were recorded under a stereomicroscope. We measured the the oocyte length for each parasitoid species using Image J software. Six individuals of *P. vindemiae* and eight *T. drosophilae* were dissected.

**1.3 Statistical analyses**

Time spent in every oviposition step and the total host handling time of *T. drosophilae* and *P. vindemiae* and their oocyte numbers were compared using Student’s t-tests. We then tested if the lifetime potential fecundity and oocyte size were significantly differed between *T. drosophilae* and *P. vindemiae* using U-Mann–Whitney tests, since the data were not normally distributed.

All statistical analyses were processed using the statistical software R version 4.2.1.

2 **Results**

**2.1** **Oviposition dynamic of two parasitoids**

The mean number of eggs produced by *T. drosophilae* per day was higher than that of *P. vindemiae* (*t*_13.28_ = -2.53, *p* = 0.025). *Trichopria drosophilae* laid 64% of eggs within five days of emergence, and the maximum daily number of eggs was 25 on the second day after emergence. By contrast, *P. vindemiae* laid 93% of eggs within seven days of emergence, and the maximum daily number of eggs was 11 on the fourth day after emergence (Figure S1). The lifetime potential fecundity of *T. drosophilae* females was twice that of *P. vindemiae* (137 ± 22.10 and 50 ± 12.93 for *T. drosophilae* and *P. vindemiae*, respectively, Mann-Whitney U tests, *w* = 84, *p* < 0.001). The ovigeny index of *T. drosophilae* (0.379 ± 0.045) was significantly higher than for *P. vindemiae* (0.159 ± 0.041, *w* = 70, *p* < 0.001).

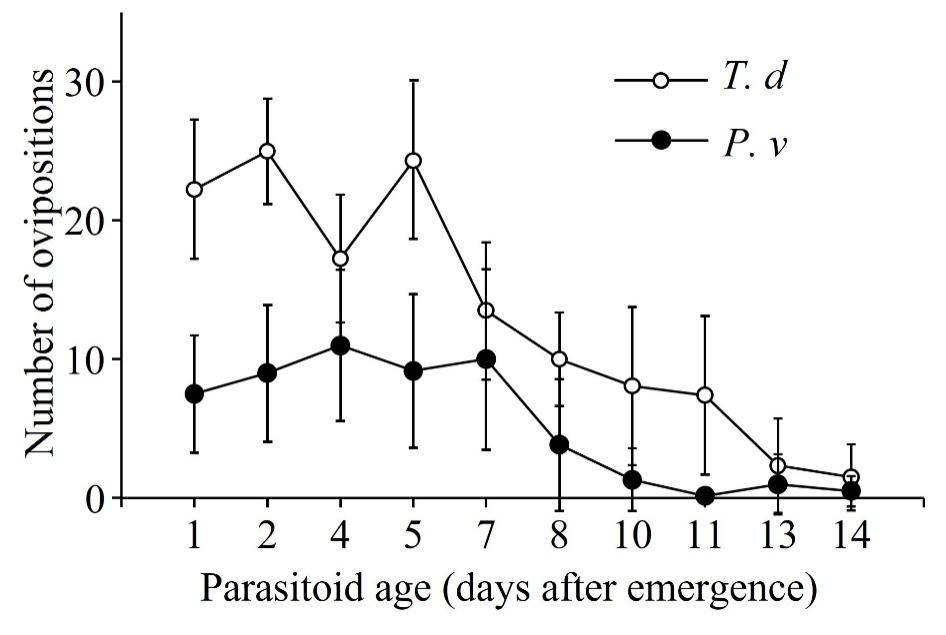

Figure S1, Number of eggs produced by *Trichopria drosophilae* (*T.d*) and *Pachycrepoideus vindemiae* (*P.v*) per day within 14 days of emergence.

**2.2 Reproductive traits of adult parasitoids**

**Host handling time**

The two parasitoid species have similar host handling steps including drumming and oviposition, and they spend more than 75% of the total handling time on the final oviposition stage (Table S1). Time spend on drilling for *T. drosophilae* was significantly longer than for *P. vindemiae* (T-tests, *t*_13.95_ = 2.60, *p* = 0.021), while no significant difference in the time taken by the other steps for the two parasitoids was detected.

Table S1 Time spend in ovipositing step for *Trichopria drosophilae* and *Pachycrepoideus vindemiae*. Mean ± se are shown, n = 8 for each species. Different letters show significant differences between the two parasitoid species (*p* < 0.05).

|  | *Trichopria drosophilae* | *Pachycrepoideus vindemiae* |
| --- | --- | --- |
| Drumming | 34.4±8.3 | 52.0±26.1 |
| Drilling | **28.8±8.1a** | **17.5±8.6b** |
| Oviposition | 197.3±50.5 | 241.0±66.3 |
| Total (seconds) | 260.4±54.6 | 310.5±66.0 |

**Oocytes number and size**

The average number of oocytes in the ovary of a newly emerged *T. drosophilae* was significantly higher than that of *P. vindemiae* (*t*_6.90_ = 17.32, *p* < 0.001). But *P. vindemiae* producing significantly larger oocytes compared with *T. drosophilae* (U-Mann–Whitney tests, *w* = 39, *p* < 0.001, Table S2).

Table S2 Number and length of oocytes from newly emerged *Trichopria drosophilae* and *Pachycrepoideus vindemiae*. Different letters indicate significant differences between the two parasitoid species (*p* < 0.05).

|  | *Trichopria drosophilae* | *Pachycrepoideus vindemiae* |
| --- | --- | --- |
| Oocyte number | 52.3±6.2a | 8.0±2.2b |
| Oocyte length (mm) | 0.5±0.05a | 0.7±0.05b |

**References：**

Jervis, M. A., Heimpel, G. E., Ferns, P. N., Harvey, J. A., & Kidd, N. A. (2001). Life‐history strategies in parasitoid wasps: a comparative analysis of ‘ovigeny’. *Journal of Animal Ecology,* 70(3), 442-458. https://doi.org/10.1046/j.1365-2656.2001.00507.x

Vayssade, C., Martel, V., Moiroux, J., Fauvergue, X., Van Alphen, J.J., & Van Baaren, J. (2012). The response of life-history traits to a new species in the community: a story of Drosophila parasitoids from the Rhône and Saône valleys. *Biological Journal of the Linnean Society,* 107(1), 153-165. https://doi.org/10.1111/j.1095-8312.2012.01918.x
